## Supplementary table 1 for "A comparison of software for analysis of rare and common short tandem repeat (STR) variation using human genome sequences from clinical and population-based samples"

|  |  | GangSTR | | | | | HipSTR | | | | | ExpansionHunter | | | | |
| --- | --- | --- | --- | --- | --- | --- | --- | --- | --- | --- | --- | --- | --- | --- | --- | --- |
| Sample | Read length/bp | Called at 30x (%) | Called at 100x (%) | Calls in both (%) | Identical by allele length (%) | Identical by allele sequence (%) | Called at 30x (%) | Called at 100x (%) | Calls in both (%) | Identical by allele length (%) | Identical by allele sequence (%) | Called at 30x (%) | Called at 100x (%) | Calls in both (%) | Identical by allele length (%) | Identical by allele sequence (%) |
| HG002 | 2x150 | 95.5 | 98.2 | 90.7 | 99.9 | NA | 91.6 | 91.5 | 88.1 | 99.6 | 99.3 | 94.8 | 99.7 | 94.8 | 98.5 | NA |
| HG003 | 2x150 | 95.4 | 98.2 | 90.7 | 99.9 | NA | 91.5 | 91.4 | 88.0 | 99.2 | 98.8 | 94.8 | 99.7 | 94.8 | 98.4 | NA |
| HG004 | 2x150 | 95.7 | 97.6 | 90.6 | 99.9 | NA | 91.2 | 90.9 | 87.7 | 99.2 | 98.8 | 99.2 | 99.7 | 99.1 | 98.4 | NA |
| HG005 | 2x250 | 86.6 | 98.2 | 85.3 | 99.9 | NA | 70.4 | 71.1 | 66.7 | 99.6 | 98.6 | 95 | 99.7 | 95 | 95.8 | NA |
| HG006 | 2x150 | 95.1 | 98.2 | 90.5 | 99.9 | NA | 91.3 | 91.3 | 87.6 | 99.1 | 98.7 | 94.8 | 99.7 | 94.8 | 98.3 | NA |
| HG007 | 2x150 | 95.6 | 97.5 | 90.4 | 99.9 | NA | 91.2 | 91.0 | 87.6 | 99.1 | 98.7 | 99.2 | 99.7 | 99.1 | 98.3 | NA |
| NA12878 | 2x150 | 95.9 | 97.6 | 90.7 | 99.9 | NA | 90.8 | 90.5 | 87.0 | 99.0 | 98.6 | 99.2 | 99.7 | 99.2 | 98.3 | NA |
| HG002 | 2x250 | 93.3 | NA | NA | NA | NA | 70.6 | NA | NA | NA | NA | 99.6 | NA | NA | NA | NA |
| HG003 | 2x250 | 93.1 | NA | NA | NA | NA | 69.6 | NA | NA | NA | NA | 99.6 | NA | NA | NA | NA |
| HG004 | 2x250 | 94.1 | NA | NA | NA | NA | 70.3 | NA | NA | NA | NA | 99.7 | NA | NA | NA | NA |

### Supplementary table 1 – Percentage of loci called in their respective STR catalogues for by GangSTR and HipSTR and ExpansionHunter
